# MTORC2 is a physiological hydrophobic motif kinase of S6 Kinase 1

**DOI:** 10.1101/2021.08.17.456612

**Authors:** Sheikh Tahir Majeed, Rabiya Majeed, Aijaz A Malik, Muhammad Afzal Zargar, Khurshid Iqbal Andrabi

## Abstract

Ribosomal protein S6 kinase 1 (S6K1), a major downstream effector molecule of mTORC1, regulates cell growth and proliferation by modulating protein translation and ribosome biogenesis. We have recently identified eIF4E as an intermediate in transducing signals from mTORC1 to S6K1 and further demonstrated that the role of mTORC1 is restricted to inducing eIF4E phosphorylation and interaction with S6K1. This interaction relieves S6K1 auto-inhibition and facilitates its hydrophobic motif (HM) phosphorylation and activation as a consequence. These observations underscore a possible involvement of mTORC1 independent kinase in mediating HM phosphorylation. Here, we report mTORC2 as an *in-vivo/*physiological HM kinase of S6K1. We show that rapamycin-resistant S6K1 truncation mutant ΔNHΔCT continues to display HM phosphorylation with selective sensitivity toward Torin-1. We also show that HM phosphorylation of wildtype S6K1and ΔNHΔCT depends on the presence of mTORC2 regulatory subunit-rictor. Furthermore, truncation mutagenesis and molecular docking analysis reveal the involvement of a conserved 19 amino acid stretch of S6K1 in mediating interaction with rictor. We finally show that deletion of the 19 amino acid region from wild type S6K1 results in loss of interaction with rictor, with a resultant loss of HM phosphorylation regardless of the presence of functional TOS motif. Our data demonstrate that mTORC2 acts as a physiological HM kinase that can activate S6K1 after its auto-inhibition is overcome by mTORC1. We, therefore, propose a novel mechanism for S6K1 regulation where mTOR complex 1 and 2 act in tandem to activate the enzyme.

## Introduction

The mechanistic target of rapamycin (mTOR), also known as the mammalian target of rapamycin, is the master regulator of cell growth and proliferation, whose deregulation is implicated in various pathological conditions including cancer, and diabetes, arthritis, and osteoporosis (1–4). mTOR forms structurally and functionally two distinct complexes called the mTOR complex 1 (mTORC1) and mTOR complex 2 (mTORC2) (5–8). mTORC1 comprises three core components viz the catalytic subunit-mTOR, the regulatory subunit-raptor, and GβL (also known as mLst8). In addition, PRAS40 and Deptor constitute the inhibitory subunits of mTORC1. On the other hand, mTORC2 also comprises mTOR and GβL. However, instead of the raptor, the regulatory component of mTORC2 is a 200 KDa protein rictor (9). The other components of mTORC2 are PRR5, Deptor, and SIN1(10). Among the two complexes, mTORC1 is extensively studied and implicated in integrating signals from growth factors and nutrients to regulate cell growth and proliferation. One of the major downstream effector molecules of mTORC1 is P70 ribosomal protein S6 kinase 1 (S6K1), which is a member of the AGC family of protein kinases (11, 12). S6K1 is regulated by multiple phosphorylation events by different kinases, which are regulated by growth factor signaling and nutrient availability (13). S6K1 exists in two isoforms (αI and αII isoforms), which are transcribed from a single gene by alternative mRNA splicing, using an alternative translational start site (14, 15). The αI isoform, containing 525 amino acid residues, has an additional 23 amino acid residue segment at N-terminus that encodes a nuclear localization motif, whereas the αII isoform, having 502 amino-acid residues, is cytoplasmic and starts at a Met residue equivalent to Met-24 in the αI isoform, and the sequences of both isoforms are identical thereafter (from here onward S6K1 refers to the larger αI isoform) (12, 16). Structural analysis of S6K1 reveals an N-terminal kinase domain and a 104-amino acid C-terminal auto-inhibitory domain (AID) and a C-terminal tail region that separates N and C termini (Fig 1b). S6K1 activation has been attributed to multiple distinctive phosphorylation events carried out by different enzymes(16–18). Among these phosphorylation events, phosphorylation at Thr412 (Thr-389 in αII isoform) in the hydrophobic motif (HM) of the tail region of the enzyme is considered a critical phosphorylation for the complete activation of S6K1.The kinase responsible for the HM site (Thr-412) phosphorylation was found to be mammalian target of rapamycin (mTOR) (19, 20). As per the prevailing view, S6K1 uses amino terminal sequence motif TOS (TOR Signaling) to interact with mTORC1 regulatory subunit raptor, that typically recruits mTORC1 substrates (21). This interaction is believed to induce S6K1 phosphorylation at Thr-412 present in the hydrophobic motif (HM) (12–14). Accordingly, loss of Thr-412 phosphorylation due to mTORC1 inhibition by rapamycin endorses the dependence of hydrophobic motif (HM) phosphorylation on the activation state of mTORC1. Contrarily however, the data demonstrating continued display of HM phosphorylation in S6K1 mutants, which harbour mutations disrupting TOS function, suggest that mTORC1 regulates S6K1 in a manner that is distinct from the one that regulates HM phosphorylation (22, 23).These observations are further supported by the recent data that demonstrates raptor is not always involved in S6K1 phosphorylation (24).Furthermore, identification of other mTORC1 substrates without a consensus TOS motif also undermine the centrality of TOS in mediating raptor binding (25–27). Notably, our recent data identifies eukaryotic translation initiation factor 4E (eIF4E) as an intermediate in transducing signals from mTORC1 onto S6K1 (28). Therein, we demonstrate that the role of mTORC1 is restricted to engaging eIF4E with S6K1-TOS motif for relieving the auto-inhibition due to carboxy terminal auto-inhibitory domain (CAD) and thereby facilitating the consequential HM phosphorylation and activation of S6K1.These observations, being consistent with our earlier data (29, 30), highlight the discordance in mTORC1 mediated HM phosphorylation and therefore suggest an alternative mechanism for the regulation of HM phosphorylation of S6K1.

**Fig 1.**
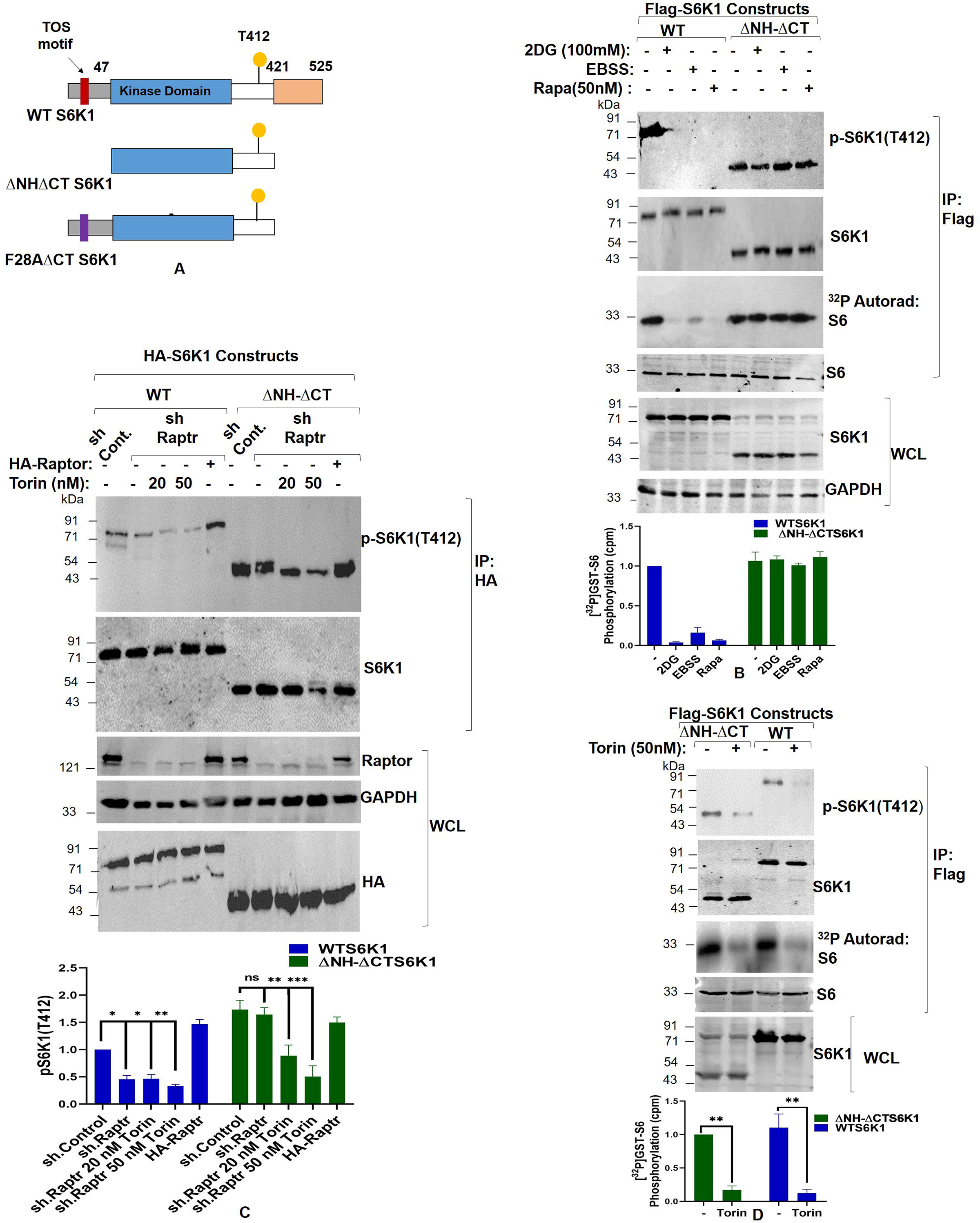
mTORC1 does not phosphorylate hydrophobic motif of S6K1. (A) S6 Kinase 1 domain structure. Representative picture demonstrates the structure full length S6K1 along with the location of TOS motif. Also shown are the truncation mutant ΔNHT2 and F28AΔCT used in the study. (B) Release of auto-inhibition renders S6K1 independent of mTORC1 regulation. Flag-WTS6K1 and Flag-ΔNHT2 S6K1 stable HEK293 cells were either lysed directly or after incubation with agents known to inhibit mTORC1 input (100 mM 2-Deoxy Glucose or 50 nM rapamycin for 30 minutes). Alternatively, mTORC1 inhibition was achieved by growing cells in amino acid free media (i.e., Earl’s balanced Salt Solution-EBSS).The lysates obtained were Flag immunoprecipitated and probed for T412 levels of S6K1. Furthermore, S6K1 kinase activity was monitored using GST-S6 as a substrate in an *in vitro* kinase assay. Quantitation represents average results of three independent experimental series. Error bars denote SEM. (C) Raptor is not involved in mediating hydrophobic motif phosphorylation of S6K1. Raptor or scrambled shRNA infected HEK293 cells were transfected with wild type S6K1 or truncation mutant ΔNHT2. Before lysis, the cells were treated with increased concentration of torin in indicated manner for 30 minutes. The lysates obtained were HA immunoprecipitated and probed for T412 levels of S6K1. Additionally, 2 μg of HA-rictor encoding plasmid was transfected in rictor shRNA cell line to rescue its knock down effect. Quantitation showingT412 phosphorylation levels represents average result of two independent experimental series normalized to the respective loading control protein content. Error bars denote SEM. p value denoted as * indicates P<0.05, ** as P< 0.005 and *** as P<0.0005 (D) S6K1 amino and carboxy termini truncation mutant ΔNHT2 is sensitive to torin. Flag-WTS6K1 and Flag-ΔNHT2 S6K1 stable HEK293 cells were either lysed directly or after 30 minutes incubation with 50nM torin. The lysates obtained were Flag immunoprecipitated and probed for T412 levels of S6K1 and S6K1 activity as described in (B).Quantitation represents average results of three independent experimental series. Error bars denote SEM. p value denoted as ** indicates P<0.005.

In this study, we report mTORC2 as a physiological HM kinase of S6K1. We identify a 19 amino acid sequence in S6K1 responsible for mediating mTORC2 dependent HM phosphorylation. We further conclude that mTORC2 mediated phosphorylation and activation of S6K1 is subservient to mTORC1 mediated release of S6K1 auto-inhibition. Our data proposes a novel mechanism describing a stepwise action of mTOR complex 1 and 2 in regulating S6K1 activation.

## Materials and Methods

### Cell lines and Culture Conditions

HEK293, HEK293T cells and NIH3T3 cells, described previously (28), and MEFs were maintained in Dulbecco’s modified Eagles medium (DMEM) supplemented with 10% (v/v) foetal bovine serum (Invitrogen), 50 μg/ml penicillin and 100 μg/ml streptomycin (Sigma-Aldrich) at 37°C with 5% CO2.

### Reagents and Antibodies

PVDF membrane (GE Healthcare/Millipore), Rapamycin (Sigma Aldrich), Torin-1 (Sigma Aldrich) and Protein G-Agarose beads (Genscript), Polyetheleneimine reagent (Polysciences, Inc.), Radioactive ATP (BRIT, Hyderabad-India). Antibody against p-S6K1 (T389/T412) was purchased from abcam, USA. Antibodies against Raptor, Rictor, S6 and Flag-tag were purchased from Cell Signaling Technologies (Beverly MA); HA-tag, myc-tag and GAPDH (Sigma-Aldrich); S6K1 (GenScript); rabbit and mouse secondary antibodies conjugated to IR Dye 800CW (LI-COR Biotechnology, Lincoln, Nebraska).

### Expression Vectors and Transfections

A pMT2-based cDNA encoding an amino-terminal HA-tagged S6K1 (αI variant) and S6K1 truncation mutant ΔNHΔCT S6K1 were gifted by Prof. Joseph Avruch, Harvard Medical School Boston, USA. Truncation mutants of S6K1 *viz* 147-421, 110-421, 91-421 and 77-421 were constructed in pKMYC mammalian expression vector. A common reverse primer 5’ GCCGAATTCCTAACTTTCAAGTACAGATGGAG3’ and specific forward primers 5’GAGGATCCATGCTGGAGGAAGTAAAGCAT3’; 5’GCGGATCCATGAAAGTAACAGGAGCAAATACT3’; 5’GCGGATCCATGTTTGAGCTACTTCGGGTACTTG3’and 5’ GCGGATCCATGACTAGTGTGAACAGAGGGCCA 3’ were used for PCR amplification of the respective mutants. cDNA HA-Raptor (#8513), HA-Rictor (#1860) were purchased from addgene. HA tagged S6K1 truncation mutant S6K1Δ91-109 was generated using primers (1) 5’Phos-AAAGTAACAGGAGCAAATACTGGGAAGATA3’ and (2)5’Phos-ACATTCTGGTCTGATTTTTTCTGGCCC3’. The mutations were verified by DNA sequence and restriction analysis. For transient transfection of cells, 1 × 10^6^ cells were plated onto a 60-mm dish 24 hr prior to transfection. 1-2 μg of Plasmid DNA along with transfection agents Lipofectamine (Invitrogen) or Polyetheleneimine, PEI (Polysciences, Inc.) were used to transfect the cells. Standard protocol was followed for cell transfection.

### Stable cell lines

Stable cell lines of HEK293 overexpressing S6K1 were generated by retroviral transduction of pWZL-Neo-Myr-Flag-RPS6KB1 expression plasmid (addgene#20624). Stable cell lines of HEK293 overexpressing S6K1 mutant ΔNHT2 were generated by retroviral transduction of pWZL-Neo-Myr-Flag-ΔNHT2 S6K1 plasmid, generated by gateway cloning (31). Briefly, the PCR amplified ΔNHT2 S6K1 was sub-cloned into TOPO adapted entry vector using pENTR/TEV/D-TOPO Cloning Kit (Invitrogen). The entry vector was then recombined with pWZL-Neo-Myr-Flag-DEST Gateway-compatible retroviral destination vector (addgene# 15300) using LR Clonase™ II Enzyme mix (Invitrogen 11791020). The infected cells were selected and maintained with neomycin (500 μg/ml).

### Gene Knock Down using shRNA

Non-target scrambled shRNA (SHC002) was purchased from Sigma Aldrich. shRNA to human raptor (plasmid#1857) and rictor (plasmid#1853) were purchased from Addgene. The preparation of shRNA infected HEK293 cells have been described previously (28).

### Immuno-precipitations and Western blotting

48 h post transfection, cells were starved overnight in serum-free DMEM. Cells were serum stimulated for 30 minutes in presence or absence of 50 nM of rapamycin or torin-1 as per the experimental requirements before lysis with ice cold NP40 lysis buffer (32). Centrifugation (micro centrifuge, 13,000 rpm at 4°C for 20 min) was carried out to remove the precipitated material to obtain a clear lysate. After clearing, the supernatant was added to 2μg each of either anti-HA, anti-Myc or anti-Flag antibodies (as per the experimental requirements) immobilized on 20 μl of a 50% slurry of protein G Agarose and rotated for 4 hours at 4°C. Immunoprecipitates were washed five times with lysis buffer. 2X Laemmli sample buffer was added to the immunoprecipitates. The samples were boiled at 100°C, centrifuged at 13,000 rpm for 2 minutes and resolved by SDS-PAGE. Proteins were transferred on PVDF membrane, probed with different antibodies at indicated concentrations and analysed using Odyssey infrared imager (LI-COR).

### In vitro kinase assay

*In Vitro* Kinase assay has been described previously (29, 33). Briefly, Proteins immobilized on either HA or Myc-beads were incubated with 1μg of the substrate and 5μCi ^32^PATP in a kinase reaction buffer containing 50mM Tris-Cl (pH 7.0), 10mM MgCl2, 0.5mM DTT, 50mM β-Glycero-phosphate, and 1mM unlabelled ATP for 15 minutes at 37°C. Reaction was stopped by adding 5X loading buffer, run on an SDS-PAGE gel. Proteins were transferred on PVDF membrane, auto-radiographed and activity determined by standard protocol.

### Molecular Docking

The amino-acid sequence of rictor from residues 971-1040, representing its stability region, which remains critical for mTORC2 function, was submitted to I-TASSER (34, 35). The protein model was subjected to energy minimization using GROMOS96 force field. Structure validation of the final model was performed using programs in NIH SAVES server (http://nihserver.mbi.ucla.edu/SAVES/), including PROCHECK, WHATIF, Verify3D and Ramachandran map. 3D model of the full length S6K1, based on the crystalline structure (PDB 4L42), was used. The modeled S6K1 and the stabilizing region of rictor were subjected to ClusPro2.0 server [2] for determining their contact interface (36). The intermolecular docking with the largest cluster of interactive residues with the lowest local energy was selected. Pymol software (The PyMOL Molecular Graphics System, Version 1.3r1 edu, Schrodinger, LLC, NY, USA) was used for building the molecular interactive protein structure models.

### Computational alanine scanning

The computational alanine scanning was used with mutations for the 19 amino acid segment of S6K1 domain. The binding energy for the mutated system (MT) and wild type (WT) system of the complex was used for the FoldX approach (37, 38). The graphical user interface for the FoldX calculations was used, which was supplemented as a plugin for the YASARA molecular graphics software. If the free energy change between mutant and wild type, denoted as ΔΔG = ΔG (MT) - ΔG(WT), is > 0, the mutation can be seen as destabilizing while ΔG < 0 suggests that the mutation will be stabilizing the protein or complex (39, 40).

### Statistical Analysis

Statistical analysis was performed using GraphPad software (Prism 8). All error bars represent SEM. For multiple group comparisons all groups from each experimental repeat were compared using 2way ANOVA. If the ANOVA test was significant (p< 0.05), Tukey’s test was performed. The asterisks denote a significant p value defined as * for P<0.05, ** for P <0.005 and *** for P<0.0005.

## RESULTS AND DISCUSSION

### MTORC1 is not a hydrophobic motif kinase of S6K1

To determine whether TOS motif is indispensable for mediating Thr-412 phosphorylation of S6K1 and whether relieving –NH2 and –COOH termini renders S6K1 active and independent of-the-regulation by mTORC1, we used previously described HEK293 cells stably expressing Flag-tagged wild type S6K1 (28). In addition, we created lines of HEK293 cells stably expressing Flag-tagged S6K1 truncation mutant ΔNHΔCT to observe their state of Thr-412 phosphorylation in response to mTORC1 inhibition. We chose ΔNHΔCT S6K1 for the purpose because it carries truncation mutations at TOS motif bearing -NH2 terminal domain and -COOH terminus auto-inhibitory domain (Fig 1a). mTORC1 inhibition was achieved either by treating the cells with glycolytic inhibitor 2 deoxy glucose (2DG) or rapamycin or by growing in Earl’s Balanced Salt Solution (EBSS) (22). As seen, ΔNHΔCT HEK293 stable cells display Thr-412 phosphorylation to an extent equivalent to their wild type counterpart that also correspond with its ability to phosphorylate GST-S6 *in vitro* (Fig 1b). Furthermore, it was interesting to observe that, unlike wildtype S6K1, ΔNHΔCT S6K1 HEK293 cells exhibit complete resistance to mTORC1 inhibition (Fig 1b).Taken together, the data indicate that mTORC1 by itself does not appear to be the HM kinase and may only serve as a priming step for the HM phosphorylation of S6K1. However, it could still be argued that ΔNHΔCT S6K1, being an unnatural variant, may support raptor recruitment co-incidentally by virtue of some unknown sequence to facilitate HM-phosphorylation. We, therefore, downregulated the expression of raptor by lentivirally transducing HEK293 cells using raptor shRNA and transfected them with HA tagged ΔNHΔCT S6K1 or wild type S6K1 to assess whether ΔNHΔCT S6K1 still sustains HM phosphorylation. Fig 1c clearly demonstrates that raptor knockdown induces loss of Thr-412 phosphorylation in wild type S6K1 only and no such loss was observed in ΔNHΔCT S6K1. Altogether, the data substantiates that the kinase other than mTORC1 was responsible for mediating Thr-412 phosphorylation in S6K1. Pertinently, the Sabatini group has reported the involvement of mTORC2 in mediating HM phosphorylation of ΔNHΔCT S6K1 (23). However, their proposition that mTORC2 mediated HM phosphorylation is a non-physiological/random event determined by the structure of S6K1 does not reconcile with the data, including ours, that highlight the redundancy of TOS motif in mediating mTORC1 specific phosphorylation of S6K1 or other bonafide mTORC1 substrates, and therefore point towards an alternative mechanism for S6K1 regulation (25–28, 33). To address this conundrum, we first ascertained the involvement of mTORC2 in mediating HM phosphorylation of ΔNHΔCTS6K1. We used HEK293 cells stably expressing Flag-tagged ΔNHΔCT-S6K1 for the purpose. Since ΔNHΔCT-S6K1 is a rapamycin resistant variant (22, 23, 28), we used torin-1, which is a specific inhibitor of mTOR kinase, for observing Thr-412 phosphorylation levels. Selective sensitivity of ΔNHΔCT S6K1 towards torin-1, that was also reflected through *in vitro* kinase assay using GST-S6 as substrate, confirmed the prospect of mTORC2 as a potential S6K1 HM kinase (Fig 1c, 1d).

### MTORC2 phosphorylates S6K1 at Thr-412 site present in the hydrophobic motif

In light of the above results that point towards the involvement of mTORC2 in mediating HM phosphorylation, we examined the state of HM phosphorylation and activation of S6K1 following knockdown of mTORC2 regulatory subunit-rictor. Accordingly, we transfected wild type HA-tagged S6K1 in rictor shRNA or, alternatively, in control shRNA infected HEK293 cells to detect any change in the magnitude of Thr-412 phosphorylation present in HM. As seen, Thr-412 phosphorylation in rictor shRNA infected HEK293 cells decreases as compared to control shRNA HEK293 cells (Fig 2A). Interestingly, this decrease in Thr-412 phosphorylation was more prominent upon treatment with mTORC1 specific inhibitor-rapamycin (Fig 2A), suggesting a combined action of mTORC1/2 in mediating T412 phosphorylation.

**Fig 2.**
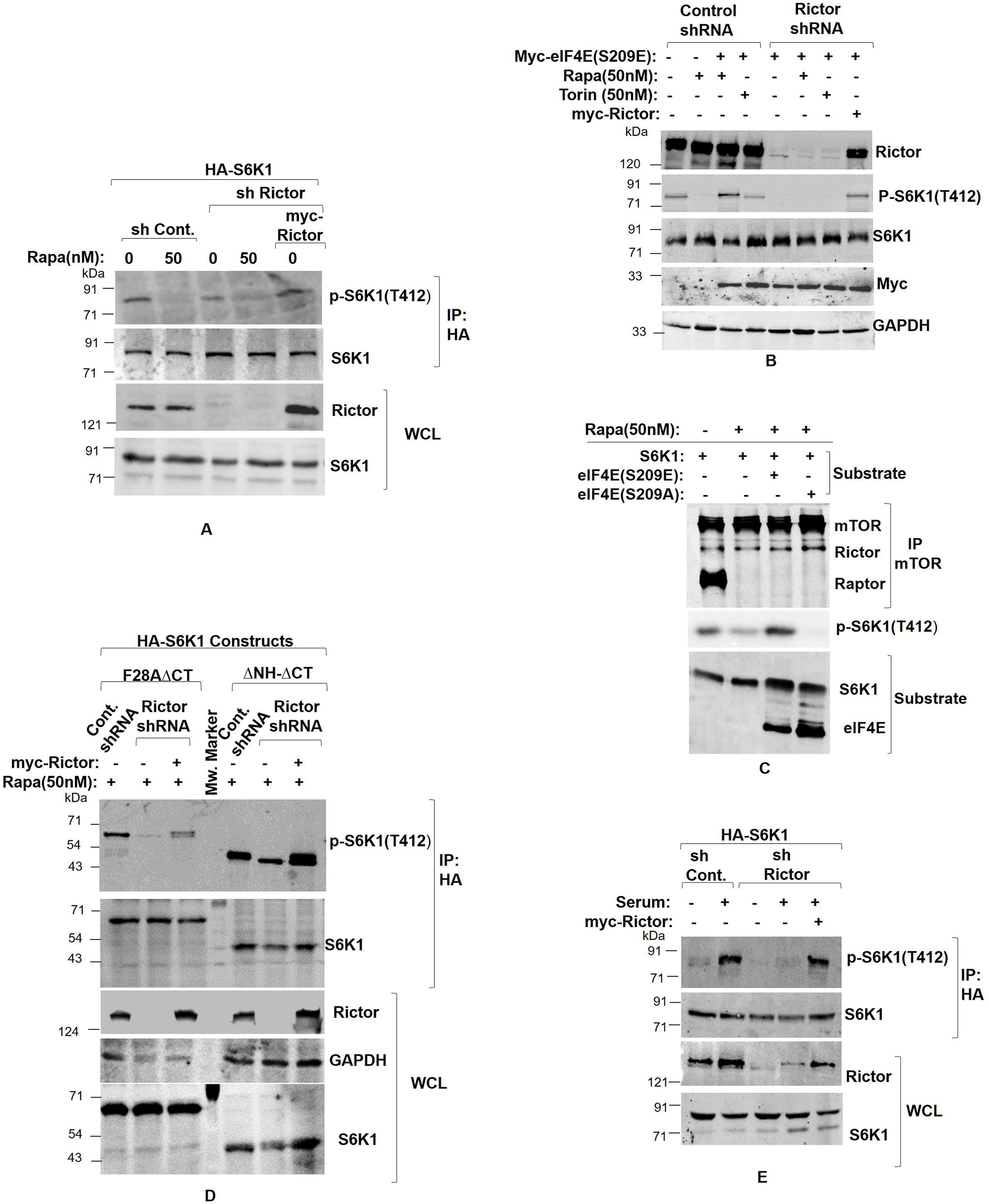
mTORC2 is an *in vivo* T412 kinase of S6K1. (A)S6K1 loses T412 phosphorylation in rictor knock down cells. HEK293 cells were infected with rictor shRNA to generate rictor knockdown cell line. Scrambled shRNA was used as control. The cells were transfected with HA tagged WT-S6K1. Before lysis, the cells were treated with increased concentration of rapamycin for 30 minutes in indicated manner. The lysates obtained were HA immunoprecipitated and probed for T412 levels of S6K1. Quantitation showingT412 phosphorylation levels represents average result of three independent experimental series normalized to their respective protein content. Error bars denote SEM. p value denoted as ** indicates P < 0.005. (B) Phosphomimetic eIF4E does not sustain T412 phosphorylation in rictor knockdown cells. Myc-tagged eIF4E was transfected in rictor or scrambled shRNA infected HEK293 cells in indicated manner. Additionally, 2 μg of myc-rictor encoding plasmid was transfected in rictor shRNA cell line to rescue its knock down effect. Prior to lysis, the cells were treated with 50nM of rapamycin or torin-1 for 30 minutes in indicated manner. The lysates obtained were probed for T412 levels. (C) Phosphomimetic eIF4E (S209E) over-rides rapamycin inhibition *in vitro*. mTOR was immunoprecipitated from rapamycin treated mouse embryonic fibroblasts (MEFs) and assessed for phosphorylating purified S6K1 in presence of purified phospho mutants of eIF4E. S6K1 and eIF4E phospho-mutants (S209A and S209E) were purified from HEK293T cells transfected with pEBG-S6K1, pEBG-eIF4ES209A and pEBG-eIF4ES209E plasmids respectively. The lysates were GST pull down, thrombin cleaved to obtain the purified substrates and used for in vitro kinase assay. The immunoblots were probed by indicated antibodies (D) HA-tagged rapamycin resistant S6K1 mutants F28AΔCTand ΔNHΔCT were transfected in rictor or scrambled shRNA infected HEK293 cells in indicated manner. Additionally, 2 μg of myc-rictor encoding plasmid was transfected in rictor shRNA cell line to rescue its knock down effect. Prior to lysis, the cells were treated with 50nM of rapamycin for 30 minutes. The lysates obtained were HA immunoprecipitated and probed for T412 levels. (E) S6K1 T412 phosphorylation does not respond to serum stimulation in rictor knock down cells. HA tagged wild type S6K1 was transfected in rictor or scrambled shRNA infected HEK293 cells in indicated manner. Additionally, 2 μg of myc-rictor encoding plasmid was transfected in rictor shRNA cell line to rescue its knock down effect. The cells were grown in serum supplemented DMEM for 48 hours and then serum starved for 12 hours. Prior to lysis, cells were serum stimulated for 30 minutes in indicated manner. The lysates were HA immunoprecipitated and probed for T412 levels.

Whilst, we have previously demonstrated that the role of mTORC1 is restricted to inducing eIF4E interaction with TOS motif of S6K1. In the same study, we show ectopic expression of phosphomimetic eIF4E overrides rapamycin mediated mTORC1 inhibition and continues to sustain S6K1 activation (28). We, therefore, explored whether phosphomimetic eIF4E could sustain S6K1 activation under rictor knockdown state. Accordingly, we transfected rictor or scrambled (control) shRNA infected HEK293 cells with phosphomimetic eIF4E (S209E) to observe S6K1 Thr-412 levels in presence of rapamycin or torin-1. As seen, control shRNA infected cells transfected with myc-eIF4E(S209E) retainThr-412 phosphorylation only in presence of rapamycin (mTORC1 inhibitor) while as torin-1 (mTORC1/C2 inhibitor) brings about significant loss of Thr-412 phosphorylation. Taken together, the data suggests the involvement of mTORC1 in priming of the enzyme and that of mTORC2 in mediating HM phosphorylation (Fig 2B). Interestingly, phosphomimetic eIF4E does not recover Thr-412 phosphorylation in rictor shRNA infected HEK293 cells. However, when co-transfected with myc-rictor, which was used to rescue rictor knockdown effect, HEK293 recovered Thr-412 phosphorylation to significant levels (Fig 2B).

We next used *in vitro* kinase assay to test if the phosphomimetic eIF4E over-rides rapamycin inhibition in similar manner as described above. Accordingly, we immunoprecipitated mTOR from rapamycin treated MEFs and assessed its potential to phosphorylate purified S6K1 in presence of purified phospho-mutants of eIF4E. S6K1 and eIF4E phospho-mutants (S209A and S209E) were purified from HEK293T cells transfected with pEBG-S6K1, pEBG-eIF4ES209A and pEBG-eIF4ES209E plasmids respectively. The lysates were GST pull down and subsequently thrombin cleaved to obtain the purified substrates. As seen, mTOR immunoprecipitates from rapamycin untreated MEF cell lines, containing both the raptor-mTOR and rictor-mTOR complexes, phosphorylate S6K1 at T412 site, while as the mTOR immunoprecipitates from rapamycin treated MEFs, containing only rictor-mTOR complexes, marginally phosphorylate S6K1 (Fig 2C). However, upon addition of the purified phosphomimetic eIF4E (S209E), the mTOR immunoprecipitates from rapamycin treated MEFs phosphorylate S6K1 significantly whereas no such phosphorylation was observed in presence of purified phosphodeficient eIF4E (S209A) (Fig 2C). Taken together, the data on one hand confirms the involvement of mTORC2 in mediating Thr-412 phosphorylation of S6K1, while on the other, it suggests that rapamycin induced mTORC1 inhibition fails to release S6K1 auto-inhibition and as a result prevents mTORC2 mediated HM phosphorylation of the enzyme. Although, the observation appears in tune with our previous finding that demonstrates the role of mTORC1 in relieving S6K1 auto-inhibition and priming it for HM phosphorylation and subsequent activation(28), we wanted to further substantiate our findings. Therefore, we used two different rapamycin resistant mutants of S6K1 viz. ΔNHΔCT and F28AΔCT for observing their state of Thr-412 phosphorylation in rictor knock down HEK293 cells(23). While ΔNHΔCT mutant is described above, F28AΔCT shares the same –COOH terminal truncation as ΔNHΔCT but bears an inactivating mutation in the TOS motif instead of the -NH2 terminal truncation (Fig 1A) (23, 30). As seen, both the mutants, despite being rapamycin resistant, exhibit significant loss of Thr-412 phosphorylation in rictor knock down state, indicating the involvement of mTORC2 in mediating Thr-412 phosphorylation (Fig 2D). The result further strengthens our view point that the loss of Thr-412 phosphorylation by rapamycin mediated mTORC1 inactivation in wild type S6K1 is due to the persistent auto-inhibition, caused by CAD, which restricts the access for mTORC2 to phosphorylate S6K1 at Thr-412 site. We also evaluated the impact of serum stimulation on endogenous Thr-412 phosphorylation levels of S6K1 in shRNA rictor knockdown NIH3T3 cells. We chose NIH3T3 cells for the purpose because the activation of endogenous S6K1 by serum in NIH3T3 cells is 2-3 fold higher than that observed in HEK293 cells (13). The results obtained suggest that serum does not induce Thr-412 phosphorylation in rictor shRNA cells as compared to control shRNA cells (Fig 2E). Altogether, the data demonstrates that Thr-412 phosphorylation is indeed mediated by mTORC2 after disinhibition or priming by mTORC1.

### A 19 amino-acid region of S6K1 mediates mTORC2 phosphorylation

Although the data presented above validates mTORC2 as a physiological Thr-412 kinase, we further wanted to ensure the specificity of mTORC2 in mediating this phosphorylation. We first argued, if mTORC2 mediated T412 phosphorylation in above discussed truncation mutants of S6K1 is a non-physiological/ random event, as proposed by the Sabatini group (23), then a smaller mutant of S6K1 should also be phosphorylated in a similar manner by mTORC2. To address this argument, we examined the sequence of ΔNHΔCT S6K1, which comprises a total of 375 amino acids starting from amino acid 47 to 421 (see Fig 1A). A careful examination of the sequence revealed conserved TOS like motif, spanning the region from amino acid 154 to 158 (Fig 3A).Therefore, we truncated 100 amino acids from -NH2 terminal end of ΔNHΔCT to generate a smaller mutant 147-421, which behaves like a TOS swap variant of S6K1 (Fig 3B). This mutant was important in two aspects, as it would tell whether the presence of TOS like motif enhances its prospect of phosphorylation by mTORC1, and whether the smaller size of the mutant better facilitates its non-physiological/random phosphorylation at Thr-412 site by mTORC2. Therefore, we transfected 147-421 S6K1 or wild type S6K1 or ΔNHΔCT S6K1 in HEK293 cells and observed the state of Thr-412 phosphorylation. Unlike wild type S6K1 and ΔNHΔCT S6K1, the mutant 147-421 failed to display any detectable phosphorylation at Thr-412 (Fig 3C), indicating redundancy of TOS motif in mediating this phosphorylation on one hand as well as dispelling the notion that mTORC2 mediated Thr-412 phosphorylation is a non-physiological event on the other. Instead, the results suggest involvement of a truncated region i.e. 47-146 in regulating Thr-412 phosphorylation. Therefore, to identify the sequence imparting Thr-412 regulation, we generated a series of smaller deletions *viz* 77-421 S6K1, 91-421 S6K1 and 110-421 S6K1 across this amino acid stretch and transfected them in HEK293 cells. Whereas the mutants 77-421 S6K1 and 91-421 S6K1 exhibit Thr-412 phosphorylation, no such phosphorylation was observed in 110-421 S6K1 mutant (Fig 3D). Also, addition of torin-1 resulted in loss of the Thr-412 phosphorylation in both 77-421 S6K1 and 91-421 S6K1 mutants (Fig 3D).Together, the data highlights the involvement of this 19 amino acid stretch, starting from amino acid 91 to 109, in imparting Thr-412 regulation. Although we furthered our attempt to examine smaller deletions i.e., 91-99 and 100-109 within this stretch, but failed to gather any information due to the unreliable expression of these mutants. Next, we deleted these 19 amino acid residues from wildtype S6K1 to generate Δ91-109 S6K1. We, then, transfected HEK293T cells with Δ91-109 S6K1 mutant to observe whether the presence of TOS bearing amino terminus and the carboxy terminus would help the mutant recover its Thr-412 phosphorylation. As seen in Fig 3E, the mutant fails to display any detectable Thr-412 phosphorylation, which was corroborated with the inability of the mutant to phosphorylate GST-S6 substrate in an *in vitro* kinase assay. Taken together, the data suggests that this 19 amino acid stretch is responsible for mediating mTORC2 dependent HM phosphorylation of S6K1.

**Fig 3.**
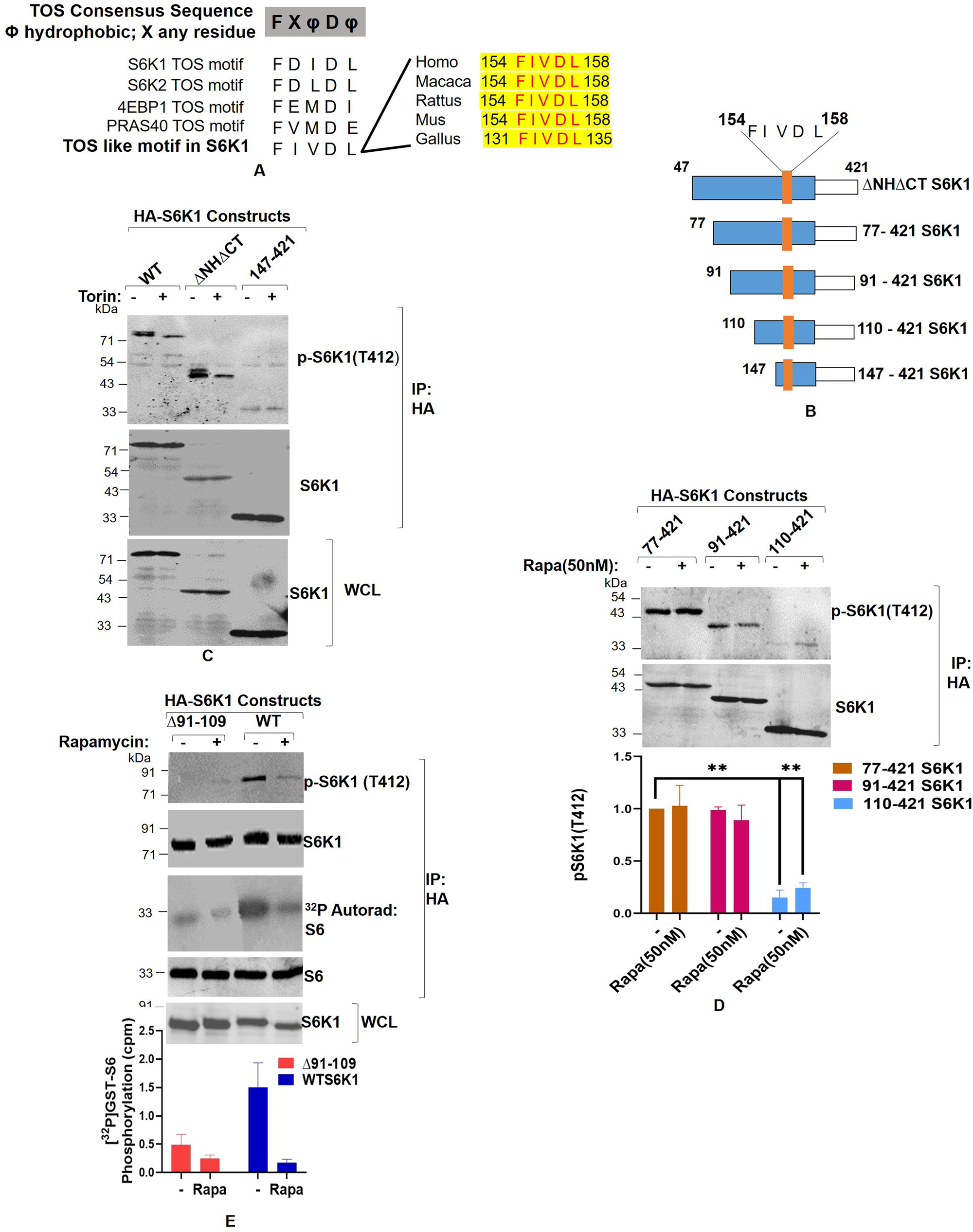
S6K1 harbors mTORC2 responsive site. (A) S6K1 harbors a TOS like motif in kinase domain. Sequence alignment shows a TOS like motif in catalytic kinase domain of S6K1 and its conserved nature across the species. (B) Structure of truncation mutants of S6K1 used in the study. (C-E) A 19 amino acid region is responsible for mediating mTORC2 dependent T412 phosphorylation of S6K1. HEK293 cells were transfected with HA tagged wild type S6K1 or truncation mutant ΔNHΔCT or myc tagged 147-421 S6K1 as indicated (C) or with myc tagged S6K1 truncation mutants 77-421, 91-421 and 110-421 (D) or with HA tagged wild type S6K1 and Δ91-109 (E). Prior to lysis, cells were treated with 50nM of torin-1 (C and D) or with 50 nM of rapamycin (E) for 30 minutes. The lysates obtained were epitope immunoprecipitated and probed for T412 levels. Furthermore, S6K1 kinase activity was monitored using GST-S6 as a substrate in an *in vitro* kinase assay (E). Quantitation represents average results of three independent experimental series. Error bars denote SEM. p value denoted as ** indicates P < 0.005.

### S6K1 interacts with rictor

We next used intermolecular docking to evaluate the interacting potential of this 19 amino acid region of S6K1 with rictor-the regulatory subunit of mTORC2. We used amino-acid sequence of rictor from residues 971-1040, referred to as the “mTORC2 stabilization region” for *in silico* studies (34). Notably, this region plays a critical role for the activation state of mTORC2 (34). We used protein prediction webserver I-TASSER to construct 3D-modeled structure of this region using. Ramachandran plots of backbone-dihedral angles φ against ψ of amino acids in rictor stabilizing domain modeled structure revealed that over 95% of the total residues are in the allowed conformational region (Supplementary Fig 1). The observation, therefore, confirms the stability of the 3D-model. We docked this 3D model with the 3D model of the full length S6K1, obtained from the crystalline structure (PDB 4L42), to determine the contact interface. The results obtained through intermolecular docking confirms the interaction between S6K1 and the mTORC2 stabilization region. Notably, S6K1 amino acid residues involved in the interaction fall within the previously described 19 amino acid region (Fig 4 A-B). Furthermore, the charged residues of S6K1 including Glu92, Arg95, Lys99, Lys104 and Arg109, within this 19 amino acid region, exhibit strong interaction potential with the stabilizing domain of rictor which was reflected by their ability to form hydrogen and ionic bonds (Fig 4C; Table1). Interestingly, the computational alanine scanning of the amino acid residues within the 19 amino acid region of S6K1 reveals that the mutation of the amino acid residues viz Glu92, Arg95, Lys99, Gly101 and Lys104 to alanine reduce the energy of S6K1 and thereby impart stability to the enzyme (Supplementary Fig 2). It is worth mentioning that except Gly101, all the amino acid residues involved in stability of S6K1 also showed strong interacting potential with the mTORC2 stabilization region (Fig 4C; Table1)

**Fig 4.**
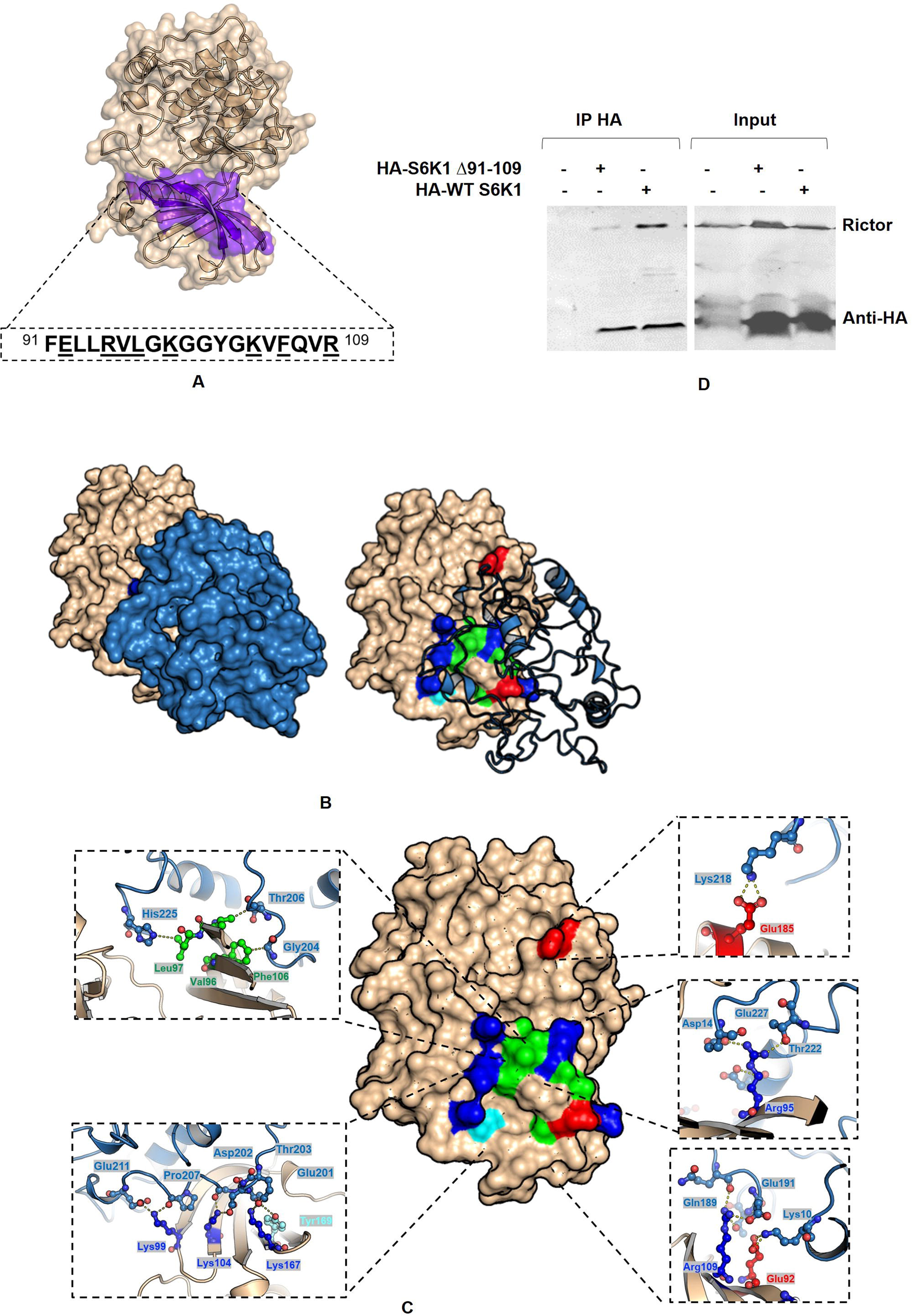
Computerized interactions between modeled S6K1 and mTORC2 stabilizing domain of rictor with focus on interactive residues. (A) S6K1 (wheat) showing the 19 amino acid segment of S6K1 (purple). The sequence of the segment is boxed and the specific residues that were predicted to form contact interfaces with stability domain of mTORC2 are underlined. (B) S6K1 (wheat) interact with stabilizing domain of mTORC2 (blue). (C) Interactive residues between S6K1 (wheat) and specific residues of stabilizing domain of mTORC2. H-bonds are indicated by yellow-colored dots. (D) HA tagged Δ91-109 S6K1 mutant alongside wild type HA-S6K1 was transfected in HEK293T cells. The lysates were HA immunoprecipitated and probed for the presence rictor.

After confirming the interaction potential of the 19 amino acid region of S6K1 with the stabilizing domain of S6K1 through *in silico* studies, we tried to confirm the results in wet lab. Accordingly, we transfected HA tagged Δ91-109 S6K1 alongside wild type S6K1 in HEK293T cells. Expectedly, immuneprecipitate of ectopically expressed wild type S6K1 co-precipitated endogenous rictor, whereas no such interaction was observed in Δ91-109 S6K1 mutant (Figure 4D).Taken together, the data confirms the interaction between S6K1 and rictor and also substantiates the role of mTORC2 as a physiological HM kinase of S6K1.

The proposal that mTORC1 is HM kinase of S6K1 while mTORC2 mediated HM phosphorylation (Thr-412) is only a non-physiological/random event determined by the structure of S6K1, does not satisfactorily reconcile with the data that demonstrate T412 phosphorylation in TOS deficient S6K1 mutants (22, 23, 28). The findings contest the preposition that interaction between raptor and S6K1-TOS motif is the basis for mTORC1 mediated T412 phosphorylation. Furthermore, absence of a consensus TOS motif in other mTORC1 substrates question the centrality of TOS in mediating mTORC1 specific substrate phosphorylation. The observation is further endorsed by our recent findings that identify eIF4E as mTORC1 substrate and an intermediate in transducing signals downstream of mTORC1 onto S6K1(28). The data, therein, demonstrates that the role of mTORC1 is restricted to engaging eIF4E with TOS motif of S6K1, required for relieving auto-inhibition and subsequently priming S6K1 for Thr-412 phosphorylation in mTORC1 independent manner (28). Based on these observations and the data presented here, mTORC2 appears to be the physiological Thr-412 kinase of S6K1. The experiments conducted in rictor shRNA infected cells, under serum-starved or serum-stimulated conditions coupled with the *in-silico* and *in-vitro* findings confirms that mTORC2 mediates Thr-412 phosphorylation, subject to the priming by mTORC1. Therefore our proposal, described in Fig 5, that S6K1 requires priming by mTORC1 dependent TOS-eIF4E engagement before it is phosphorylated at Thr-412 by mTORC2 addresses the shortcomings that otherwise discredit mTORC2 as a physiological Thr-412 kinase. Notably, deletion analysis reveals 19 amino acid stretch of S6K1 responsible for mediating Thr-

**Fig 5.**
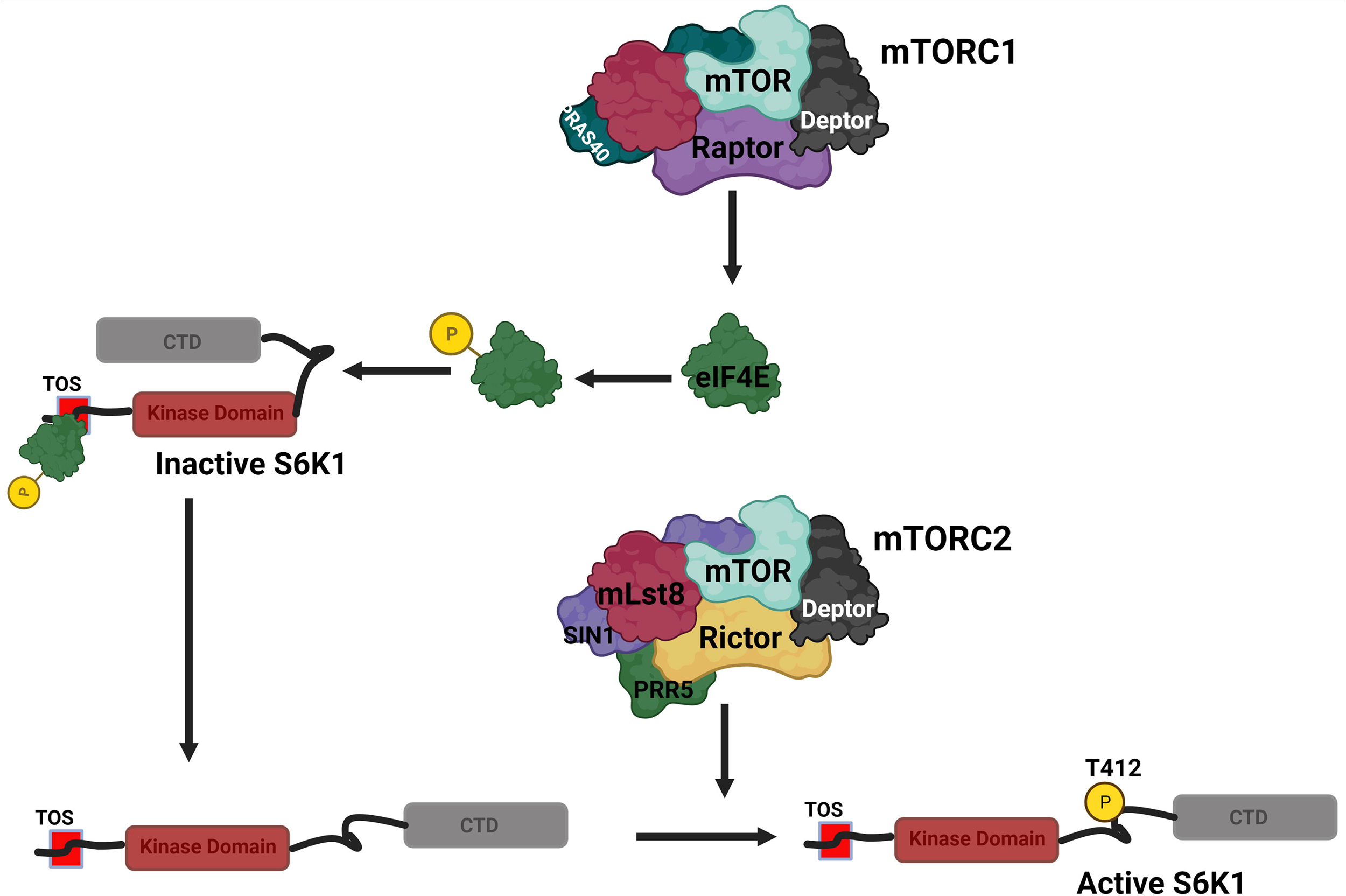
S6K1 activation model. Illustration describing a two-step activation mechanism for S6K1. Step 1. Active mTORC1 induces eIF4E to interact with TOS motif of S6K1 for releasing the S6K1 auto-inhibition and thereby exposing the critical Thr-412 site in HM for phosphorylation and subsequent activation (described previously (28)). Step2. MTORC2 phosphorylates S6K1 at Thr-412 site in HM and leads to complete activation of the enzyme.

412 phosphorylation by a torin-1 sensitive kinase and therefore associates physiological specificity with mTORC2 mediated Thr-412 phosphorylation. Furthermore, inability of the Δ91-109 S6K1 mutant, deleted of the 19 amino acid stretch, to interact with rictor and the resultant loss of its Thr-412 phosphorylation despite the presence of a consensus TOS motif, unequivocally implicates the region in mediating mTORC2 dependent T412 phosphorylation.

## Acknowledgements

We are thankful to Dr. Joseph Avruch from Harvard Medical School, Boston-MA, USA for providing cDNA clones of wildtype S6K1 and its truncation mutant ΔNHΔCT. DS Kothari Postdoctoral fellowship from University Grants Commission (UGC-India) (201920-BL/18– 19/0544) in favour of S.T.M and Senior research Fellowship from Indian Council of Medical Research (ICMR-India) (2019-5079 CMB-BMS) in favour of RM are duly acknowledged. Research and Innovation grant to KIA from Ministry of Education-Govt. of India under RUSA component 2.0 and University Grants Commission CPEPA (2–5/2016(NS/PE)) are duly acknowledged.

## Author Contributions

S.T.M. designed and performed the experiments, generated S6K1 stable cell lines, constructed S6K1 mutants, analysed and interpreted the data and wrote the manuscript. R.M. performed experiments, constructed rictor, raptor knockdown cell lines. A.M. performed the in-silico analysis. M.A.Z. supervised the project. K.I.A conceived and supervised the project, designed the experiments, interpreted the data and wrote the manuscript.

