## Supplementary figures and images for "MTORC2 is a physiological hydrophobic motif kinase of S6 Kinase 1"

### Supplementary fig 1

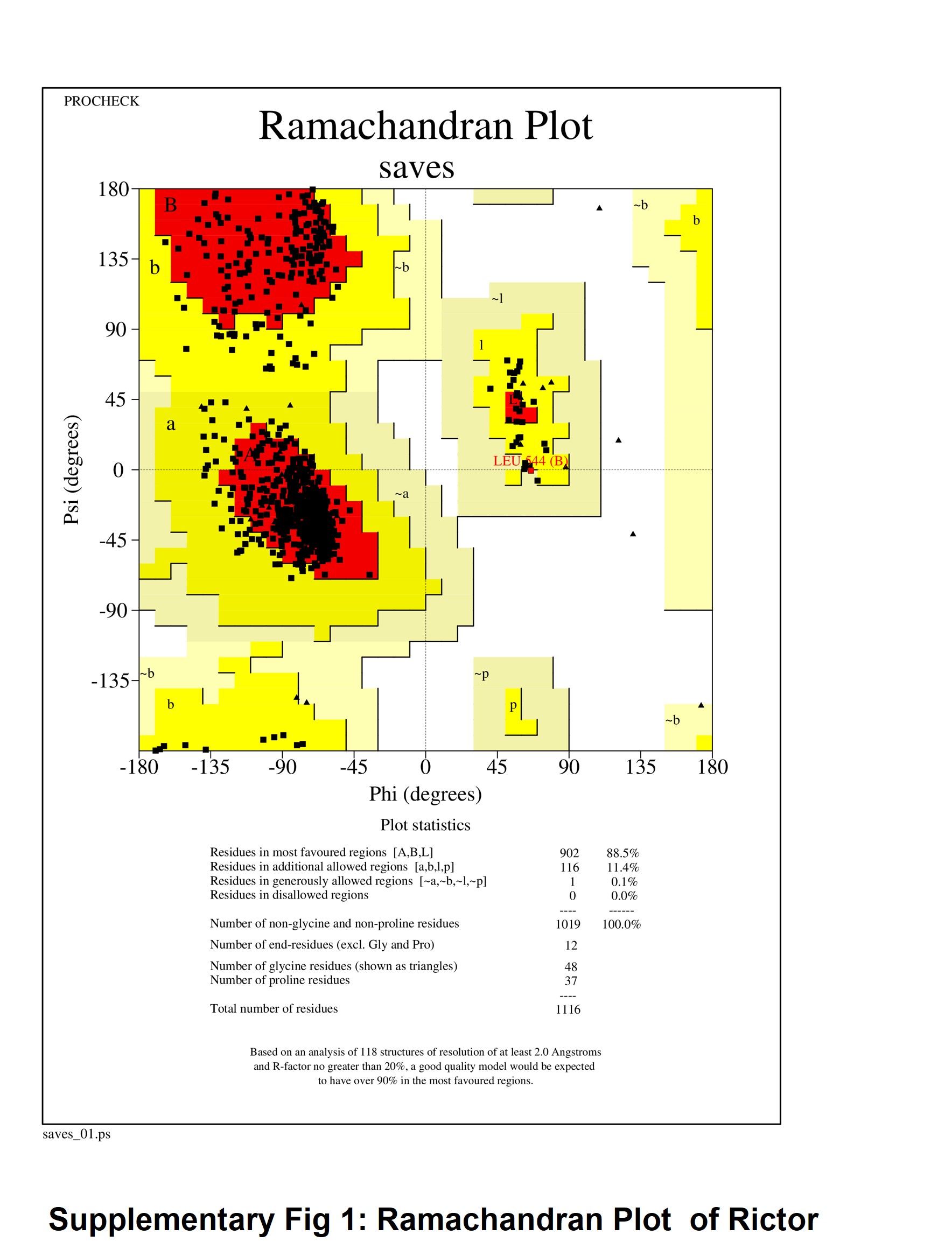

### Supplementary fig 2

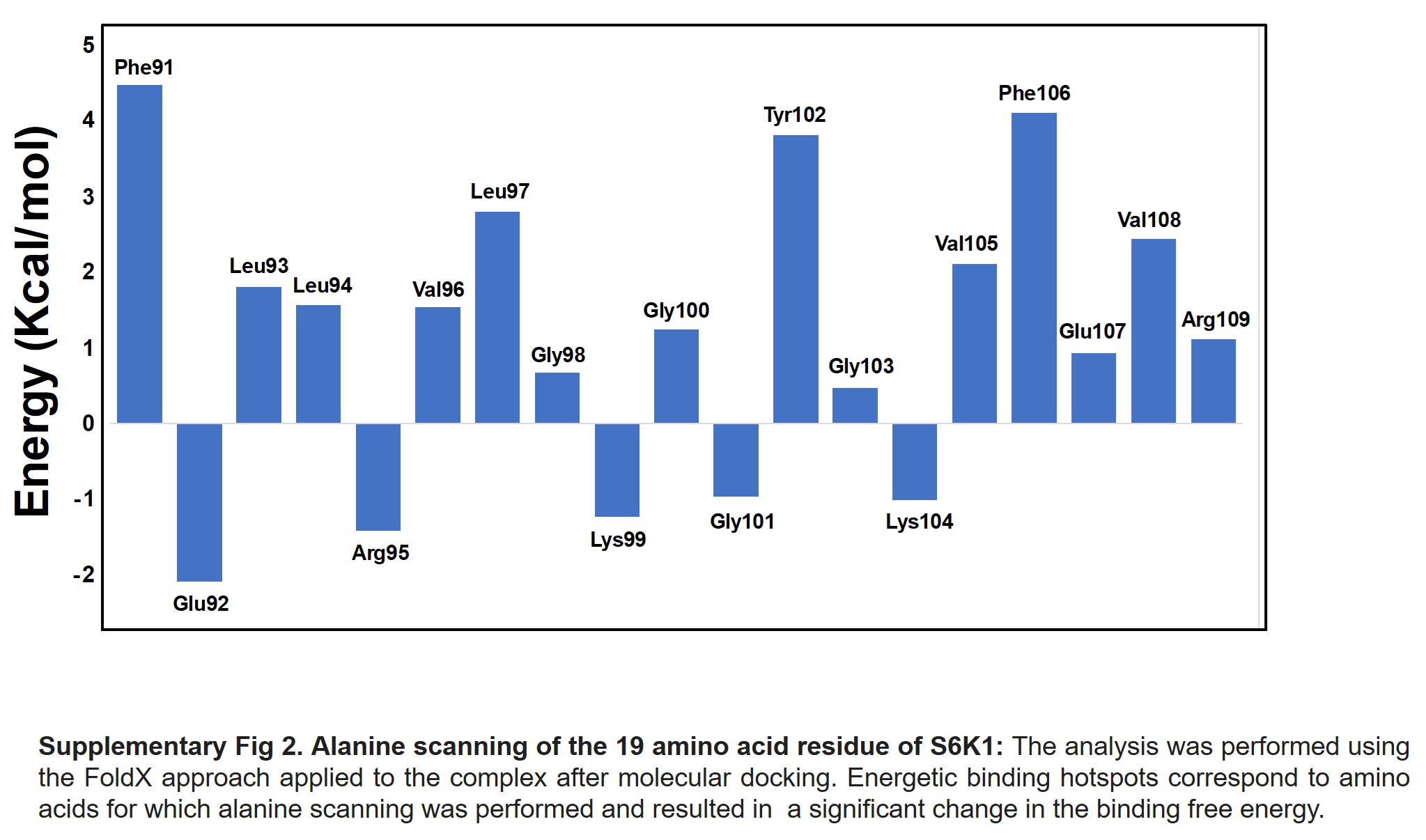
